## Supplementary material for "Freezing responses during prolonged threat memory retrieval reflect trait-like anxiety endophenotypes in female and male inbred mice": Box 1

**Anxiety** – a spectrum of approach-avoidance behaviours elicited by a potential danger/uncertainty/goal conflict

**Anxiety trait** – an intrinsic conflict sensitivity, a consistent bias towards approach or avoidance across all uncertain situations/contexts determined by (epi)genetic factors

**Anxiety state** – a transient defensive behavioural pattern activated by conflicting interests of the subject; a distinct concept from fear – a defensive behavioural pattern activated by an immediate threat

**Classical (Pavlovian) conditioning** – a procedure through which an initially neutral stimulus (conditioned stimulus, CS) is paired with a biologically potent stimulus (unconditioned stimulus, US)

**Auditory aversive conditioning (AAC)** – a procedure through which an initially neutral auditory stimulus is paired with a noxious stimulus

**Predictive value** – a certainty of one event predicting another

**Freezing** – cessation of all movements except respiratory

**Phasic freezing** – freezing occurring with the onset of the conditioned stimulus, reflects associative memory

**Sustained (tonic) freezing** – prolonged freezing occurring in some individuals, reflects a fear-potentiated anxiety state

**Phasic responders** – individuals with only phasic freezing component above baseline

**Sustained responders** – individuals with phasic and sustained freezing components above baseline
